# Fine mapping of *Ur-3*, a historically important rust resistance locus in common bean

**DOI:** 10.1101/078782

**Authors:** O.P. Hurtado-Gonzales, G. Valentini, T.A.S Gilio, A.M. Martins, Q. Song, M.A. Pastor-Corrales

**Author notes:** **Corresponding author:** Marcial A. Pastor-Corrales, USDA-ARS, Soybean Genomics and Improvement Laboratory, 10300 Baltimore Ave, Building 006, Room 118, Beltsville, MD 20705.

## Abstract

Bean rust is a devastating disease of common bean in the Americas and Africa. The historically important *Ur-3* gene confers resistance to many races of the highly variable bean rust pathogen that overcome all known rust resistance genes. Existing molecular markers tagging *Ur-3* for use in marker assisted selection produce false results. We described here the fine mapping of *Ur-3* for the development of highly accurate markers linked to this gene. An F_2_ population from Pinto 114 × Aurora was evaluated for its reaction to four different races of the bean rust pathogen. A bulked segregant analysis using the SNP chip BARCBEAN6K_3 positioned the approximate location of the *Ur-3* locus to the lower arm of chromosome Pv11. Specific SSR and SNP markers and haplotype analysis of 18 sequenced bean lines led to position the *Ur-3* locus to a 46.5 Kb genomic region. We discovered a KASP marker, SS68 that was tightly linked to the *Ur-3* locus. Validation of SS68 on a panel of 130 diverse common bean lines and varieties containing all known rust resistance genes revealed that it was highly accurate producing no false results. The SS68 marker will be of great value to pyramid *Ur-3* with other rust resistance genes. It will also reduce significantly time and labor associated with the current phenotypic detection of *Ur-3*. This is the first utilization of fine mapping to discover markers linked to a rust resistance in common bean.

## Introduction

The common bean (*Phaseolus vulgaris* L.) includes dry and snap beans. The dry edible bean is the most important pulse in the diet of humans throughout the world, especially in Latin America and Africa, where dry beans are the main daily source of protein, complex carbohydrates, fiber, and micronutrients (Broughton et al. 2003).

A myriad of biotic and abiotic factors constrain common bean production in the world. Among these, bean rust is a devastating disease that results in significant losses of seed yield in dry beans and pod quality in snap beans (Stavely and Pastor-Corrales 1989; Liebenberg and Pretorius 2010).

The bean rust disease is caused by the biotrophic basidiomycete fungus *Uromyces appendiculatus*, an obligate parasite of common bean. This pathogen has a complex life cycle with five distinct spore stages and three different nuclear conditions, which are indicative of the capacity of this pathogen for genetic recombination (Groth and Mogen 1978; McMillan et al. 2003). Many published reports reveal the rich virulence diversity of *U. appendiculatus* with scores of races (virulence phenotypes) identified around the world (Groth and Roelfs 1982; Mmbaga and Stavely 1988; Stavely and Pastor-Corrales 1989; Liebenberg 2003; Araya et al. 2004; Arunga et al. 2012; Acevedo et al. 2012). More than 90 races from the United States, Africa, Asia, and from other countries of the Americas have been characterized and maintained by the USDA-ARS Bean Project at the Beltsville Agricultural Research Center (Stavely 1984; Mmbaga and Stavely 1988; Stavely et al. 1989; Pastor-Corrales 2001).

Genetic resistance is the most cost-effective strategy to manage the bean rust disease.. Rust resistance in common bean is conditioned by single and dominant genes identified by the *Ur-* symbol (Kelly et al. 1996). To date, ten genes have been named and tagged, mostly with RAPD or SCAR molecular markers (Miklas et al. 2002). Some of these genes (*Ur-3*, *Ur-5*, *Ur-7*, *Ur-11*, and *Ur-14*) were found on common beans of the Mesoamerican gene pool, while the others (*Ur-4*, *Ur-6*, *Ur-9, Ur-12*, *and Ur-13*) were from common beans of the Andean gene pool (Augustin et al. 1972; Ballantyne 1978; Stavely 1984, 1990; Grafton et al. 1985; Finke et al. 1986; Jung et al. 1998; Liebenberg and Pretorius 2004; Souza et al. 2011).

The *Ur-3* gene present in the Mesoamerican white-seeded common bean Aurora was reported by Ballantyne in 1978. Since then, this gene has been used extensively as the source of rust resistance in a large number of dry bean varieties from various market classes of the United States, as well as in fresh market and processing snap beans (Brick et al. 2011; Urrea et al. 2009; Kelly et al. 1994; Osorno et al. 2010; Pastor-Corrales et al. 2007; Stavely et al. 1992, Beaver et al, 2015). *Ur-3* has also been used as a source of rust resistance in dry bean varieties of South Africa (Liebenberg et al. 2005). In addition, *Ur-3* has been reported in the literature as the subject of different studies including genetics (Grafton et al. 1985; Kalavacharla et al. 2000), molecular markers and gene tagging (Haley et al. 1994). The *Ur-3* gene has also been reported as occurring in Mesoamerican common bean cultivars Mexico 235, Ecuador 299, NEP 2, and 51052, that in addition to*Ur-3* also contain one or two additional rust resistance genes (Stavely et al. 1989; Miklas et al. 2000; Hurtado-Gonzales et al. 2016).

The *Ur-3* gene confers resistance to 55 of 94 races of the bean rust pathogen maintained at Beltsville, MD, USA (Pastor-Corrales et al. 2001). More importantly, *Ur-3* confers resistance to many races that overcome the resistance of all named rust resistance genes in common bean. For example, *Ur-3* is resistant to race 108, the only race known to overcome the broad spectrum resistance of the *Ur-11* gene present in PI 181996 and PI 190078, and of the *Ur-14* gene present in Ouro Negro (Stavely 1998; Alzate-Marin et al. 2004). The *Ur-3* gene also complements the broad spectrum rust resistance in accessions PI 151385, PI 151388, PI 151395, and PI 151396, which are also susceptible to race 108. Similarly, *Ur-3* confers resistance to race 84, the only known race that overcomes the rust resistance in PI 260418 (Pastor-Corrales 2005). In addition, *Ur-3* confers resistance to many races that overcome the *Ur-4*, *Ur-5*, *Ur-6*, *Ur-7*, *Ur-9*, *Ur-12*, and *Ur-13* genes. Although *Ur-3* is not resistant to all races of Mesoamerican origin, this gene confers resistance to most races of *U. appendiculatus* of Andean origin; that is, races isolated from common beans of the Andean gene pool. Thus, *Ur-3* is a critical component of gene pyramiding of common bean cultivars with broad resistance to rust. The information above provides strong evidence of the historical importance and current relevance of *Ur-3* for breeding dry and snap beans with broad and durable resistance to rust in the United States and other nations (Stavely 2000; Pastor-Corrales et al. 2001).

The resistant reaction of *Ur-3* gene to *U. appendiculatus* is initially characterized by the production of small water-soaked chlorotic spots that subsequently become well-defined necrotic spots without sporulation. This resistant phenotype is classified as grade 2 or 2,2^+^ and it is known as the hypersensitive reaction (HR) in the bean rust grading scale (Stavely et al. 1989; Stavely 1998).

The *Ur-3* gene has been mapped on chromosome Pv11 of the common bean genome (Stavely 1998; Miklas et al. 2002). Inheritance of resistance and phenotypic data revealed that the *Ur-3* gene was very closely linked to *Ur-11* on the terminal position of chromosome Pv11 (Kelly et al. 1996). The close proximity between these two genes led to the naming of the rust resistance gene in PI 181996 as *Ur-3^2^* (Kelly et al. 1996). However, later reports demonstrated the independence of *Ur-3* and *Ur-*3^2^ and revealed that these two genes were linked in repulsion and different from each other (Stavely 1998). Thus, *Ur-3^2^* was renamed *Ur-11* (Stavely 1998). The close proximity of *Ur-3* to *Ur-11* may be one of the main reasons why it has been difficult to find DNA markers that are specific the *Ur-3* gene. There are other named rust resistance genes on Pv11 (*Ur-6, Ur-7,* and *Ur-11*), as well as two other unnamed genes (*Ur-Dorado 53* and *Ur-BAC 6*), albeit, these genes are not as closely linked to *Ur-3* as *Ur-11* is (Miklas et al. 2002; Kelly et al. 2003).

Four specific races of the bean rust pathogen have been reported as phenotypic markers that effectively identify rust resistance genes; race 53 identifies *Ur-3*, race 49 for *Ur-4*, race 47 for *Ur-6*, and race 67 identifies *Ur-11* (Stavely 2000; Pastor-Corrales and Stavely 2002). These races identify the presence of these genes alone or in combinations with other rust resistance genes. However, the phenotypic identification of these rust resistance genes is laborious, time consuming, and currently only performed at the Bean Project at Beltsville. Moreover, the detection of multiple rust resistance genes in common bean using phenotypic markers is also often complicated by the presence of epistasis between rust resistance genes (Miklas et al. 1993; Pastor-Corrales and Stavely 2002). Furthermore, the current molecular markers (mostly RAPD and SCAR markers) linked to rust resistance genes in common bean that were reported almost two decades ago, yield false positive and negative results, as is the case with the currently available RAPD (OK14_620_) and SCAR (SK14) markers linked to the *Ur-3* locus (Nemchinova and Stavely 1998; Haley et al. 1994; Stavely 2000).

Several factors contributed to the false positive and false negative results when using the current molecular markers. Among these factors is the fact that some molecular markers were not sufficiently close to the gene of interest. Another constrain was the close proximity among rust resistance genes, as is the case between the *Ur-3* and *Ur-11* genes. Additionally, until recently the lack of a reference genome for common bean hindered the development of highly specific DNA markers. The publication of the common bean reference genome in 2014 (Schmutz et al. 2014) along with the development of high-throughput genotyping technologies for common bean are making possible the identification of more effective molecular markers.

The objective of this study was to develop highly effective molecular markers for the detection of the historically important and widely used *Ur-3* rust resistance gene. To accomplish this, we fine mapped the genomic region containing the *Ur-3* gene. We used a combination of high-throughput single nucleotide polymorphism (SNP) genotyping, bulked segregant analysis (BSA), and local association of the phenotype and genotype of a diverse set of 18 common bean lines. This is the first research that combines various novel technologies to fine map a bean rust resistance gene in common bean that results in the identification of a highly effective molecular marker linked to *Ur-3*.

## Material and Methods

### Population development and phenotypic evaluation of the bean rust disease

A total of 129 F_2_ plants were derived from the cross Pinto 114 x Aurora. Both are dry beans of the Mesoamerican pool of common bean, where Pinto 114 was the susceptible parent and Aurora was the resistant parent containing the *Ur-3* gene. All F_2_ plants, parents, and control cultivars were grown in 12.7-cm-diameter pots containing two plants per pot. The primary (unifoliate) leaves of bean plants were inoculated about seven days after seeding, when the primary leaves were about 35-65% expanded. To prepare the rust inocula, suspensions of frozen urediniospores of various races of *U. appendiculatus* were placed in a 25 ml solution of cold tap water and 0.01 % Tween 20 in a 250-ml Erlenmeyer flask. The spore solutions were prepared with a concentration of 2×10^4^ urediniospores.mL^−1^. All 129 F_2_ plants were inoculated with races 41, 53, 84 and 108 of *U. appendiculatus*. The following cultivars with known rust resistance genes were included in the inoculation as internal controls of successful rust inoculation: Early Gallatin (*Ur-4*), Golden Gate Wax (*Ur-6*), and PI 181996 (*Ur-11*) (Table 1). The F_2_ plants were inoculated using a cotton swab to apply the spore solution of each of the races on the abaxial side of the primarily leaves. After inoculation, the plants were transferred to a mist chamber (20 ± 1ºC and relative humidity >95%) for 18 hours under darkness. After this period, the plants were transferred to the greenhouse. Visible rust symptoms were observed on susceptible plants about 10-12 days after inoculation (dai).

**Table 1.** Reaction of the bean cultivars used in this study with races 41, 53, 84, and 108 of *Uromyces appendiculatus*.

| Cultivar | Ur gene | Races of *U. appendiculatus* |  |  |  |
| --- | --- | --- | --- | --- | --- |
|  |  | 41 | 53 | 84 | 108 |
| Pinto 114 | - | 5,4 | 5,4 | 5,4 | 5,4,6 |
| Aurora | Ur-3 | 2 | 2+ | 2+ | 2,2+ |
| Early Gallatin | Ur-4 | 4,5 | 4,5 | 4,5 | 2+ |
| Golden Gate Wax | Ur-6 | 3,f2 | 4,5 | 3,f2 | 4,5 |
| PI 181996 | Ur-11 | f2 | f2 | f2 | 5,6 |

Standard bean rust grading scale: 1 = no visible symptoms; 2, 2+ = Necrotic spots without sporulation; 3 = Tiny uredinia (sporulating pustules) less than 0.3mm in diameter; f2 = faint and tiny chlorotic spots; 4 = Medium uredinia, 0.3-0.5mm in diameter; 5 = Large uredinia, 0.5-0.8 mm in diameter, 6 = Very large uredinia, larger than 0.8mm in diameter. Reactions 2, 3, f2 are considered Resistant, and 4, 5, 6 are considered Susceptible.

The F_2_ population and parents were evaluated for their rust phenotype about 12-14 dai using a six-grade scale (Stavely and Pastor-Corrales 1989), where: 1 = no visible rust symptoms; 2 = necrotic or chlorotic spots without sporulation, less than 0.3 mm in diameter (hypersensitive reaction, HR); 2+ = necrotic spots without sporulation, 0.3-1.0 mm in diameter; 2++ = necrotic spots without sporulation, 1.0-3.0 mm in diameter; 2+++ = necrotic spots larger than 3.0 mm in diameter; 3 = uredinia (sporulating pustules) less than 0.3 mm in diameter; 4 = uredinia 0.3-0.5 mm in diameter; 5 = uredinia 0.5-0.8 mm in diameter; 6 = uredinia larger than 0.8 mm in diameter. Plants with grades 2 and 3 were classified as resistant, whereas those with rust grades of 4, 5, and 6 were classified as susceptible. Thereafter, the F_2_ plants were maintained in the greenhouse to produce F_2:3_ families by selfing. A total 281 F_3_ plants from twelve selected F_2:3_ families were inoculated with race 53 of *U. appendiculatus*. These families were inoculated using an Air Brush-Depot compressor, model TC-20, and a Iwata Airbrush, Revolution BCR, by applying the spore solution (concentration of 2×10^4^.mL^-1^) of race 53 on the abaxial side of the leaves. After spraying, plants were treated similarly to the F_2_ plants, described above.

A summary of the bean rust phenotype on differential bean cultivars with all races of the rust pathogen used during this study is presented in Table S1.

### Bulk Segregant Analysis and SNP assay

Newly trifoliate leaves from each of the F_2_ plants were collected and total genomic DNA was isolated using DNeasy 96 Plant Kit (Qiagen, CA) according to manufacturer’s instructions. Based on the rust reaction of each of the F_2_ plants, three susceptible bulks were prepared. Each bulk consisted of DNA from eight F_2_ susceptible plants, to ensure that heterozygous resistant plants were not included in the bulks. These bulks were used for bulk segregant analysis (BSA) for identification of markers potentially associated with the *Ur-3* gene (Michelmore et al. 1991). The DNA from susceptible bulks and two samples from each parent were analyzed with 5398 single nucleotide polymorphisms (SNP) markers on the Illumina BeadChip BARCBEAN6K_3 following the Infinium HD Assay Ultra Protocol (Illumina, Inc. San Diego, CA). The results obtained on the BeadChip were visualized by fluorescence intensity using the Illumina BeadArray Reader and alleles were called using Illumina GenomeStudio V2011.1 (Illumina, San Diego, CA). Allele calls were visually inspected and errors in allele calling were corrected manually. SNPs were considered associated with the *Ur-3* locus when they were polymorphic between the Pinto 114 (susceptible) and Aurora (resistant) parents and the three susceptible bulks were homozygous and clustered tightly with the susceptible parent Pinto 114.

### Developing and evaluating simple sequence repeat markers (SSRs) linked to *Ur-3*

The sequence fragments containing SNPs associated with the *Ur-3* locus were aligned to the common bean reference genome using Standalone Megablast (Morgulis et al. 2008) to identify the scaffolds in the reference genome. Scaffolds were downloaded at the Phytozome website (https://phytozome.jgi.doe.gov/pz/portal.html), DOE, JGI (Department of Energy, Joint Genome Institute). The scaffolds were screened for the presence of simple sequence repeat (SSR) markers. Procedures for SSR identification, SSR screening, and primer design were previously described by Song et al. 2010.

The polymorphism and quality of the SSR markers were first tested using DNA from the Pinto 114 (S) and Aurora (R) parents. Polymorphic SSR markers were then used to analyze the DNA of the F_2_ population from the Pinto 114 × Aurora cross. Polymerase chain reaction (PCR) was performed with 5 ng of genomic DNA, 0.25 μM of forward and reverse primers, 1 X PCR Buffer (200 mM Tris-HCl (pH 8.0), 500 mM KCl, 2mM each dNTP, 10% glycerol, 15 mM MgCl_2_, 20 ng/µL of single-stranded binding protein, 0.1 unit of Taq DNA polymerase. The PCR thermocycling parameters were: 3 min at 92°C and 38 cycles of 50 s at 90°C, 45 s at 58°C and 45 s at 72°C followed by a 5 min extraction at 72°C and hold at 10°C. PCR products were resolved on 2-3% agarose gels (Agarose SFR, Amresco) prepared with TBE 1X buffer (Tris-borate-EDTA) and stained with 1μg.mL^-1^ ethidium bromide.

### Developing and testing KASP markers

A subset of SNPs positively associated with *Ur-3* found using BSA were selected for genotyping the F_2_ population from Pinto 114 × Aurora using Kompetitive Allele Specific PCR (KASP) assays. Additional SNPs used for KASP genotyping were retrieved from SNP chip tables found in Song et al (2015). KASP primers were designed using the PrimerExpress software and KASP reactions were conducted following the manufacturer’s instructions. The 10 μL reaction contained 5 μL of 2X premade KASP master mix (LGC, Middlesex, UK), 0.14 μL of primers mix (Sigma-Aldrich, St. Louis, USA), and 20–40 ng of genomic DNA. PCR parameters were as described by LGC on standard thermocycling machines using white semi-skirted polypropylene 0.2 ml 96-well PCR plates (USA Scientific) and sealed with Microseal^®^B (Bio-Rad, Hercules, CA). After PCR amplification was completed, PCR plates were scanned using the Mx3000P qPCR machine (Agilent, St Clara, CA) and allele calls for each genotype were obtained initially using the MxPro software (Agilent, St Clara, CA) or using the Klustercaller software (LGC, Middlesex, UK).

### Construction of genetic linkage map around the *Ur-3* locus

The genetic distance between the SSRs, KASPs and the *Ur-3* locus in the F_2_ population (129 plants) was estimated using the software JoinMap 4.0 (Van Ooijen 2006). Defaults settings of a Regression Mapping algorithm based on Kosambi map function were attributed to define linkage order and distances in centimorgans (cM). A minimum likelihood of odds (LOD) ≥3.0 and a maximum distance of ≤50 cM were used to test linkages among markers.

### Fine-mapping of the *Ur-3* locus in F_3_ plants using KASP markers

F_2:3_ families were selected based on the recombination between *Ur-3* and the molecular markers (SSRs and KASPs) found in the F_2_ population. A total of ten F_2:3_ families were selected for screening with KASP markers SS4 and SS6 flanking the *Ur-3* locus. One homozygous resistant family and one susceptible family were evaluated as internal controls. The number of plants per family varied from 22 to 32 according to the availability of seeds. A total of 281 F_3_ plants were inoculated with race 53 of *U. appendiculatus* as described above. DNA from the F_3_ plants was isolated according to Lamour and Finley (2006) and were genotyped with KASP markers SS4 and SS6. F_3_ plants showing recombination between markers SS4 and SS6 were selected for additional genotyping with newly designed KASP markers in order to narrow the region containing the *Ur-3* locus.

### Haplotype analysis of the *Ur-3* locus

Haplotype analysis was performed in the genomic region flanked by the SS4 and SS6 KASP markers. These two markers flanked a region of 470,487 bp on Pv11. Eighteen diverse bean cultivars including C20, Matterhorn, Stampede, T39, Sierra, Red Hawk, Jalo EEP 558, Michelite, UC White, Kardinal, Laker, Cornell 49242, BAT 93, Buckskin, Fiero, Lark, UI 906 and CELRK were sequenced by Song et al. (2015) and used for the haplotype analysis. These lines were also inoculated with races 49, 53, 67 and 108 of *U. appendiculatus* as previously described. The sequence variants in the targeted genomic region of the 18 cultivars and their phenotypes were used to identify haplotypes associated with the resistance and susceptibility for *Ur-3*. All SNPs identified between KASP markers SS4 and SS6 were handled using Microsoft Excel and haplotypes were identified by visual inspection. At least two KASP markers were designed for each of the observed haplotypes. Whenever feasible, SNP markers were located every 10 Kb across the 470,487 bp genomic region. When KASP markers were polymorphic between the Pinto 114 (*ur-3*) and Aurora (*Ur-3*) parents, they were used to genotype F_3_ plants with recombination between the markers SS4 and SS6.

### Validation of the markers linked to the *Ur-3* locus

A panel of 130 diverse cultivars bean lines and varieties that included all rust resistance genes in common bean were genotyped using KASP markers tightly linked with *Ur-3*. This was performed with the purpose of generating accurate *Ur-3* markers useful in marker-assisted selection. In this panel, some cultivars had the *Ur-3* gene alone, other cultivars had *Ur-3* combined with other rust resistance genes, while others did not have any reported rust resistance genes. The cultivars in the panel were phenotyped before or during the course of this study with multiple races of the bean rust pathogen including race 53, the phenotypic marker for the *Ur-3* gene.

### Data Availability

All data described in this manuscript related to bean rust phenotypes, Pinto 114 x Aurora F_2_ genetic map, F_3_ fine mapping population, and haplotype analysis are available in Table S1, Table S2, Table S3, Table S4, Table S5, Table S6, and Table S7.

## Results

### Inheritance of rust resistance in common bean Aurora

A total of 129 F_2_ plants from the Pinto 114 × Aurora cross were evaluated for their reaction to races 41, 53, 84 and 108 of *U. appendiculatus* (Table S2). Aurora was resistant to all four races with a reaction that was characterized by necrotic spots without sporulation (grades 2, 2^+^). Pinto 114 was susceptible to the same four races with a reaction characterized by large uredinia (grade 4, 5, and 6). Based on the reaction to all four races, the inheritance of rust resistance study of the 129 F_2_ plants exhibited a segregation equal to 101 resistant (R) and 28 susceptible plants (S), fitting a ratio of 3R:1S (χ^2^=0.747, P value = 0.38), confirming that the rust resistance in Aurora was conferred by the single and dominant *Ur-3* gene (Table S2).

### BSA and SNP genotyping using BARCBEAN6K_3 BeadChip

Based on the BSA, 28 SNPs were associated with *Ur-3* (Table 2). The alleles of these SNPs could separate the susceptible Pinto 114 and the three susceptible bulks from the resistant Aurora parent. According to the genetic linkage map created by Song et al. (2015), these 28 SNPs were distributed from 72.3 to 76.2 cM on the lower end of the common bean chromosome Pv11. The physical location of the associated 28 SNPs was between 46,437,627 bp (ss715647455) and 48,784,158 bp (ss715641910), a region spanning a total of 2.1 Mbps (Table 2).

**Table 2.** Positive single nucleotide polymorphism (SNP) markers associated with *Ur-3* locus in the common bean linkage group Pv11. The selected SNPs segregate between the parents and *Ur-3* susceptible bulks from Pinto 114 × Aurora cross. 1=genetic position based on Song et al 2015 map.

| NCBI ssID | BARCBEAN6K_3 SNP id | Physical position<br>Pvul V1.0 | SR F<sub>2</sub> linkage map<br>position (cM)<sup>1</sup> |
| --- | --- | --- | --- |
| ss715647455 | sc00206ln407767_400194_T_G_143587659 | 46437627 | 72.337 |
| ss715639564 | sc00206ln407767_348254_C_T_143535719 | 46490018 |  |
| ss715647451 | sc00206ln407767_168453_G_A_143355918 | 46667862 | 72.512 |
| ss715647773 | sc00273ln341540_84911_C_A_168094735 | 46939681 | 73.921 |
| ss715647765 | sc00273ln341540_130419_A_G_168140243 | 46982186 | 73.921 |
| ss715647770 | sc00273ln341540_233731_C_A_168243555 | 47083906 | - |
| ss715640322 | sc00733ln158243_112892_C_T_274461500 | 47289130 | - |
| ss715649250 | sc00733ln158243_110962_C_T_274459570 | 47291059 | - |
| ss715649254 | sc00733ln158243_81656_T_G_274430264 | 47320724 | 74.735 |
| ss715649249 | sc00733ln158243_10639_C_T_274359247 | 47390755 | 75.07 |
| ss715649251 | sc00733ln158243_2983_T_C_274351591 | 47398413 | 75.07 |
| ss715649719 | sc00992ln119304_109010_C_T_310044208 | 47431965 | 75.07 |
| ss715648098 | sc00346ln293441_287396_T_C_191559002 | 47746437 | 75.684 |
| ss715648096 | sc00346ln293441_269534_C_T_191541140 | 47768651 | 75.07 |
| ss715648093 | sc00346ln293441_239954_T_C_191511560 | 47800050 | 75.07 |
| ss715649910 | sc01089ln106922_89228_T_C_320994162 | 48163156 | 75.222 |
| ss715640836 | sc01089ln106922_30683_A_G_320935617 | 48221254 | 75.222 |
| ss715648349 | sc00418ln255472_36179_T_C_211051790 | 48547014 | 76.258 |
| ss715648350 | sc00418ln255472_49970_A_G_211065581 | 48560374 | 76.258 |
| ss715648351 | sc00418ln255472_79205_G_A_211094816 | 48588580 | - |
| ss715648352 | sc00418ln255472_96331_T_G_211111942 | 48605710 | - |
| ss715648342 | sc00418ln255472_105778_T_G_211121389 | 48614962 | - |
| ss715648343 | sc00418ln255472_112102_G_T_211127713 | 48621286 | 76.258 |
| ss715648344 | sc00418ln255472_133412_T_C_211149023 | 48640040 | - |
| ss715648345 | sc00418ln255472_153753_G_A_211169364 | 48660384 | - |
| ss715648346 | sc00418ln255472_173904_T_C_211189515 | 48680290 | 76.258 |
| ss715650748 | sc01832ln56221_26620_C_T_378717205 | 48780038 | 76.258 |
| ss715641910 | sc01832ln56221_30739_G_A_378721324 | 48784158 | 76.258 |

### Mapping of the *Ur-3* gene

The large portion of the genomic region containing SNPs associated with the *Ur-3* rust resistance gene was targeted for SSR development. A total of 48 SSR markers located between 46,266,888 and 48,664,905 bp on Pv11 were developed. Thirteen of the 48 SSRs markers were polymorphic between the parents Pinto 114 (S) and Aurora (R) parents (Table 3). These markers, which showed unequivocal allele separation in agarose gel, were used to map the *Ur-3* locus in the F_2_ population Pinto 114 × Aurora. Linkage analysis positioned the *Ur-3* locus between markers BARCPVSSR14001 (46,535,562 bp) and BARCPVSSR14082 (47,291,606 bp), a 756,044 bp genomic region (data not shown). In addition, four positively associated SNPs from the BSA and two SNPs (retrieved from Song et al 2015) nearby the SSRs flanking the *Ur-3* locus were selected and converted into KASP markers (Table 4). Five KASP markers showed clear separation of the three clusters (2 homozygous and 1 heterozygous) and were polymorphic between the Pinto 114 and Aurora parents. These KASP markers were used to refine the *Ur-3* gene map. Linkage analysis in the F_2_ population genotyped with 13 SSRs and the five KASP markers showed that *Ur-3* was flanked by KASP marker SS5 and SSR marker BARCPVSSR14007 (Figure 1, Table S3). The distance of the *Ur-3* locus to both markers was 0.2 cM (Figure 1).

**Figure 1.**
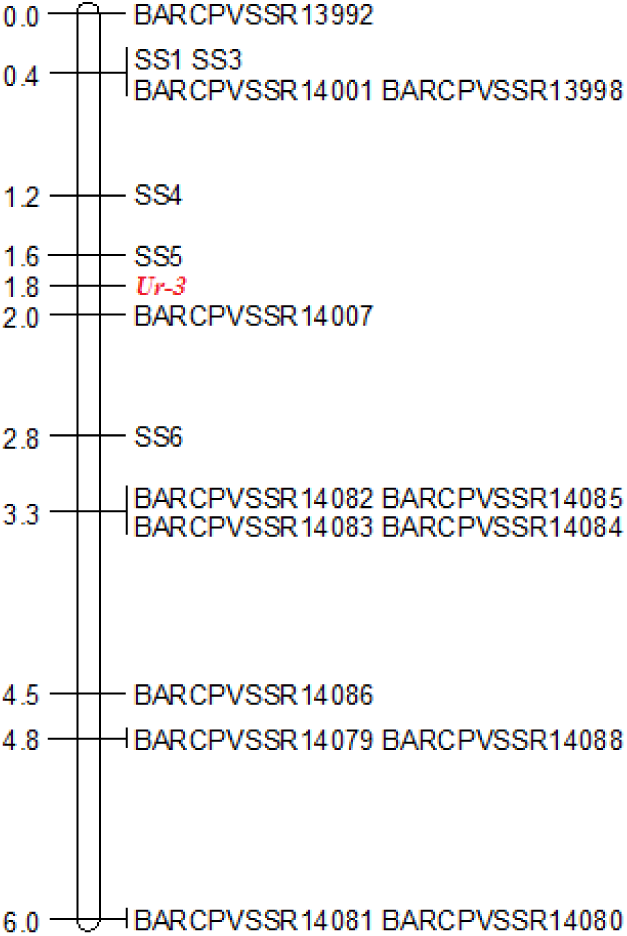
Genetic map of common bean linkage group Pv11 containing the *Ur-3* locus and the simple sequence repeats (SSRs) and single nucleotide polymorphism (SNPs) markers. KASP marker SS5 and SSR marker BARCPBSSR14007 are both linked to Ur-3 at a distance of 0.2 cM. Genetic map generated using the Kosambi’s mapping function from 129 F_2_ plants derived from Pinto 114 × Aurora cross.

**Table 3.**
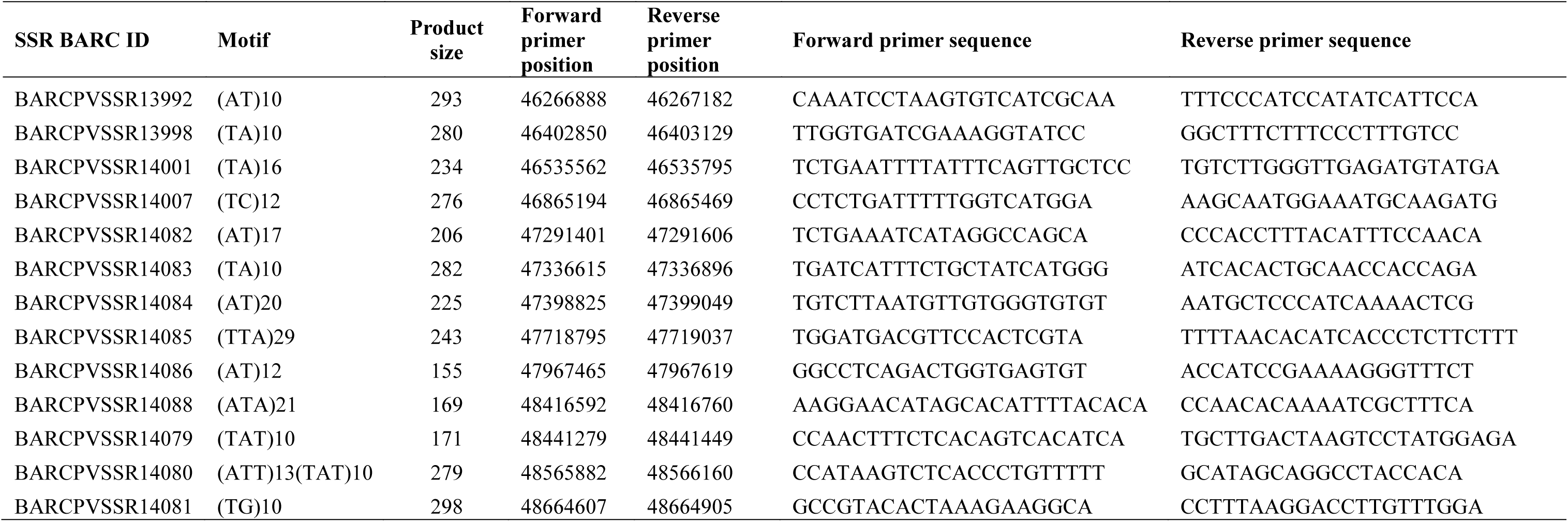
Simple sequence repeats (SSR) marker ID, motif, forward and reverse primer position n version V1.0 of the reference genome of *P. vulgaris* and primer sequences. The SSRs were ed to map the *Ur-3* locus present in Aurora cultivar.

**Table 4.**
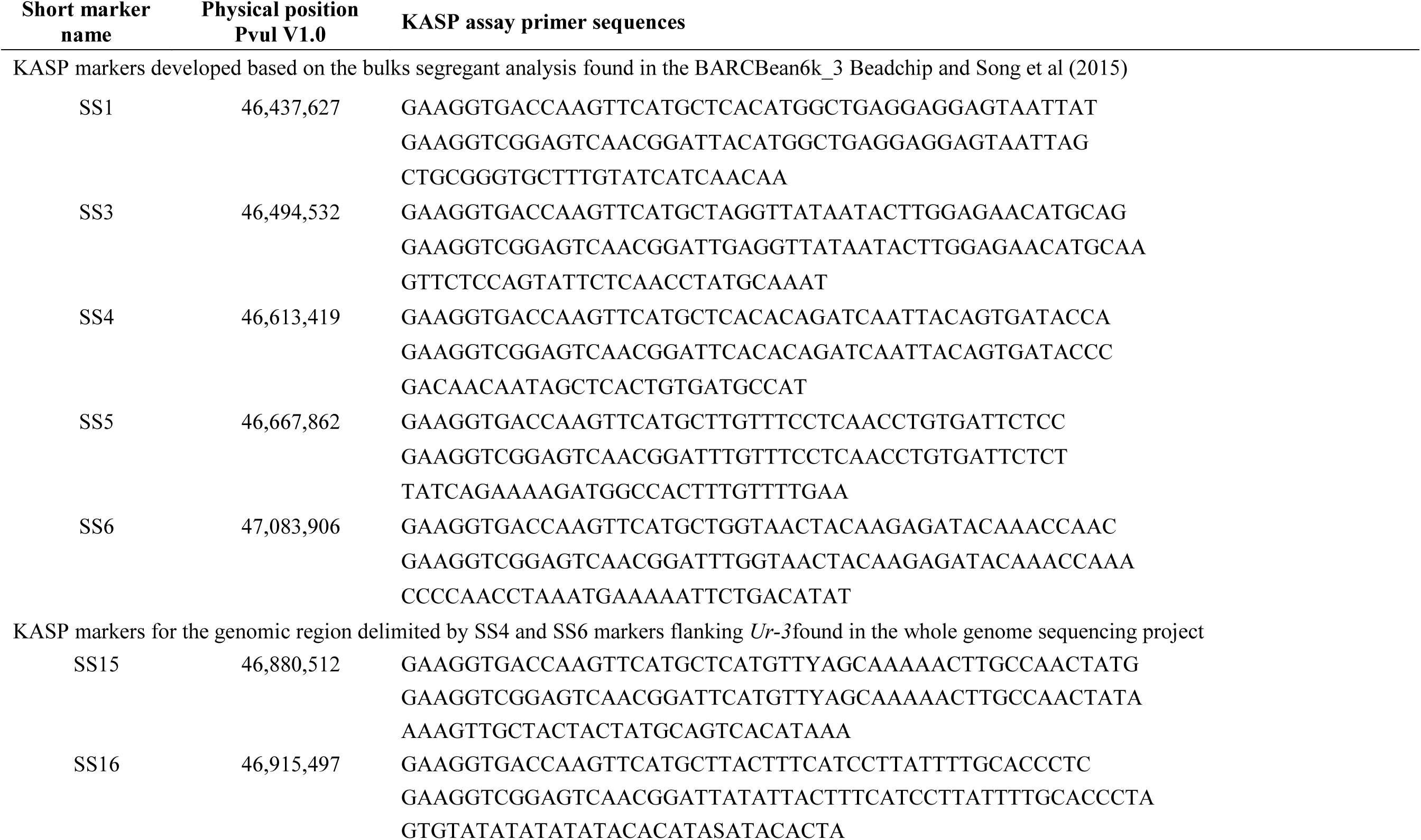
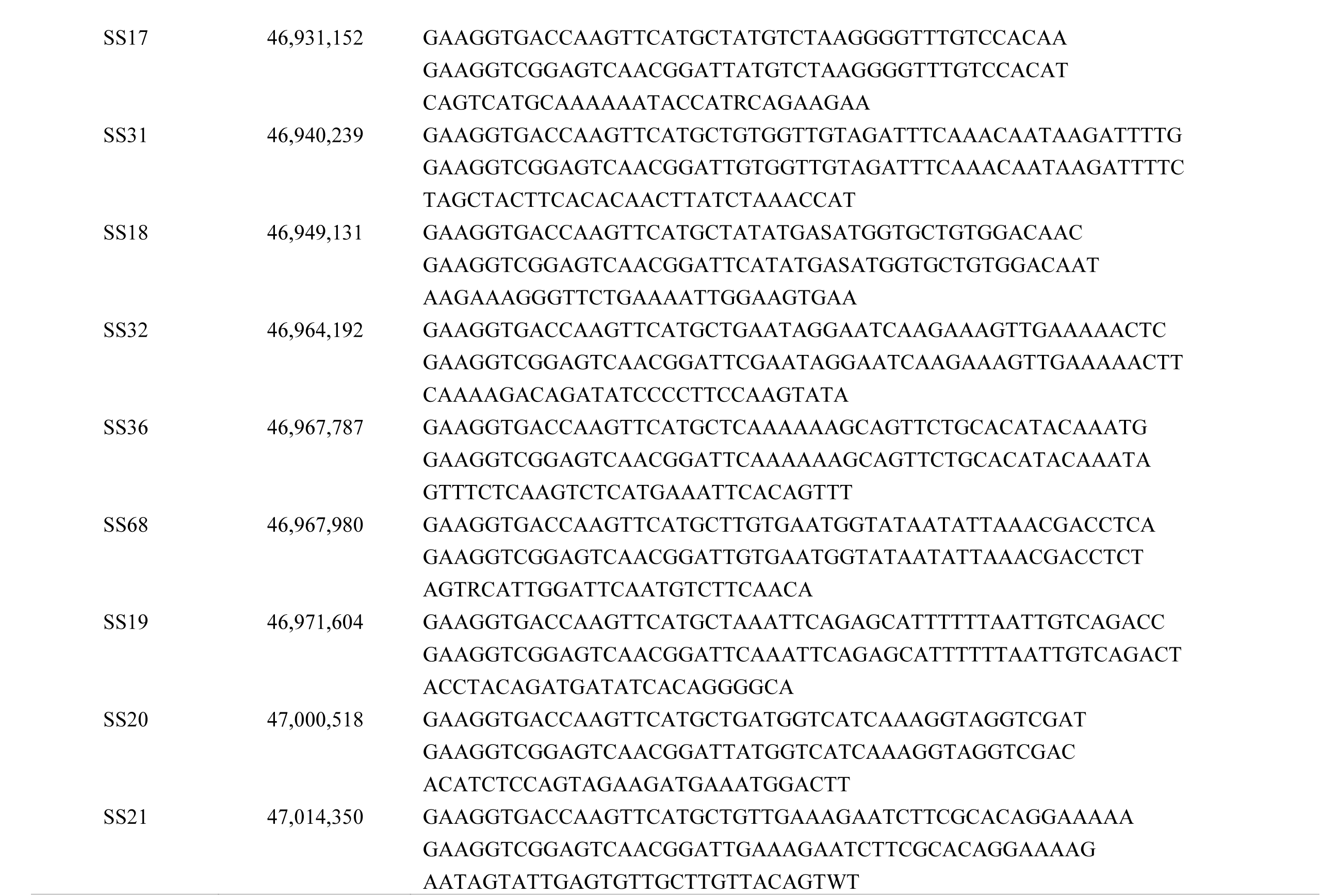
Physical position and primer sequences of KASP markers associated with *Ur-3.* KASP arkers were used to genotype F_2_ mapping population and F_2:3_ families for fine mapping.

### Analysis of recombination in F_3_ and *Ur-3* haplotype identification

KASP markers SS4 and SS6 were mapped at 0.6 and 1.0 cM from the *Ur-3* locus, respectively (Figure 1), in a 470,487 bp (470 Kb) genomic region of chromosome Pv11, from 46,613,419 to 47,083,906. These markers were chosen to genotype 12 selected F_2:3_ families from the cross Pinto 114 × Aurora. Among the 12 families, four were derived from recombinant F_2_ plants between KASP markers SS4 and SS6, six families were heterozygous between markers SS4 and SS6 flanking *Ur-3*, and two families were used as internal controls: one homozygous resistant and the other homozygous susceptible. In addition, these twelve families (281 F_3_ plants) were inoculated with race 53 of *U. appendiculatus*. Genotyping the 281 F_3_ plants resulted in 87 F_3_ plants with recombination events between the SS4 and SS6 KASP markers (Table S4). These 87 F_3_ plants were selected for subsequent fine mapping analysis with additional KASP markers (Table 4). SS5 (ss715647451 at position 46,667,862) was the only KASP marker derived from the BeanChip that was located between SS4 and SS6; thus, SS5 was also used to genotype the recombinant 87 F_3_ plants.

We then mined the sequence data (SNPs) from 18 common bean lines (Song et al. 2015) to search for additional SNPs between SS4 and SS6. Based on the whole genome sequence of the 18 common bean lines, approximately 6000 SNPs and small indels were found between SS4 and SS6 (Table S5). These SNPs were grouped into ten major haplotypes (Table 5). Each of these haplotypes were then tagged with one or two KASP markers and were examined for their polymorphism between Pinto 114 (*ur-3*), Aurora (*Ur-3*), Mexico 235 (*Ur-3+*), and PI 181996 (*Ur-11*). The KASP markers polymorphic between the Pinto 114 and Aurora parents were tested on the set of 87 F_3_ recombinant plants identified previously with KASP markers SS4 and SS6. Analysis of the 87 F_3_ recombinant plants positioned the *Ur-3* gene between KASP markers SS17 and SS21, in a genomic region of 83,198 bp (Table S7). Concurrently, a specific haplotype for *Ur-3* was identified based on the reaction of the 18 sequenced cultivars to race 53 of *U. appendiculatus.* Only the cultivars C20, Matterhorn, Stampede, T39, and Sierra had a resistant phenotype (hypersensitive response) to races 53 and 108 indicating that these cultivars have the *Ur-3* gene (Table S7). The final genotyping analysis on the 87 recombinant plants mapped *Ur-3* between KASP markers SS36 and SS21, in a specific genomic region of 46,563 bp, ranging from 46,967,787 to 47,014,350 bp of Pv11 (Table 6). Two F_3_ plants, one resistant and the other susceptible, had the same recombination breakpoint, demonstrating that the *Ur-3* gene was located in the region flanked by SS36 and SS21 (Table 6).

**Table 5.** Major haplotypes (columns) identified between SNP markers SS4 and SS6 using SNP calls from 18 sequenced bean lines (Song et al. 2015). The haplotype associated with the *Ur-3* locus is revealed by marker name SS17 at position 46,931,152. H= Heterozygous, *= missing data, Ref=Reference genome, S=Susceptible, R=Resistant

| Bean Genotype | Marker Class | Race | Marker name | SS4 | SS5 | SS15 | SS16 | SS17 | SS18 | SS19 | SS20 | SS21 | SS6 |
| --- | --- | --- | --- | --- | --- | --- | --- | --- | --- | --- | --- | --- | --- |
|  |  |  | Reaction to Race 53 | 46613419 | 46667862 | 46880512 | 46915497 | 46931152 | 46949131 | 46971604 | 47000518 | 47014350 | 47083906 |
| G19833 (Ref) |  |  | S | T | C | C | C | T | C | G | T | T | C |
| Cal Early | Light Red Kidney | Nueva Granada | S | T | C | C | * | T | C | G | T | C | A |
| Red Hawk | Dark Red Kidney | Nueva Granada | S | T | C | C | A | T | C | G | T | C | A |
| Fiero | Dark Red Kidney | Nueva Granada | S | T | C | C | A | T | C | G | T | C | A |
| Lark | Light Red Kidney | Nueva Granada | S | T | C | C | A | T | C | G | T | C | A |
| Kardinal | Light Red Kidney | Nueva Granada | S | T | C | C | A | T | C | G | T | C | A |
| BAT 93 | Tan | Mesoamerica | S | T | C | T | A | T | C | G | * | * | * |
| UC White | White Kidney | Nueva Granada | S | T | C | T | A | T | C | G | T | C | H |
| Jalo EEP 558 | Canário | Peru | S | T | C | * | C | T | C | G | T | T | A |
| UI 906 | Black | Mesoamerica | S | T | C | T | A | T | * | A | C | T | C |
| Michelite | Navy | Mesoamerica | S | G | H | C | C | T | T | A | C | T | C |
| Cornell 49-242 | Black | Mesoamerica | S | T | C | C | C | T | T | A | C | T | * |
| Laker | Navy | Mesoamerica | S | T | H | T | C | T | C | A | C | T | * |
| Buckskin | Pinto | Durango | S | T | C | T | C | T | T | A | C | C | C |
| T39 | Black | Mesoamerica | R | G | T | T | * | * | T | A | C | * | A |
| Sierra | Pinto | Durango | R | G | T | T | C | A | T | A | C | T | A |
| Matterhorn | Great Northern | Durango | R | H | C | T | C | A | * | A | C | T | A |
| Stampede | Pinto | Durango | R | G | C | T | C | A | T | A | C | T | A |
| C20 | Navy | Mesoamerica | R | H | H | T | C | A | T | A | C | T | A |

**Table 6.**
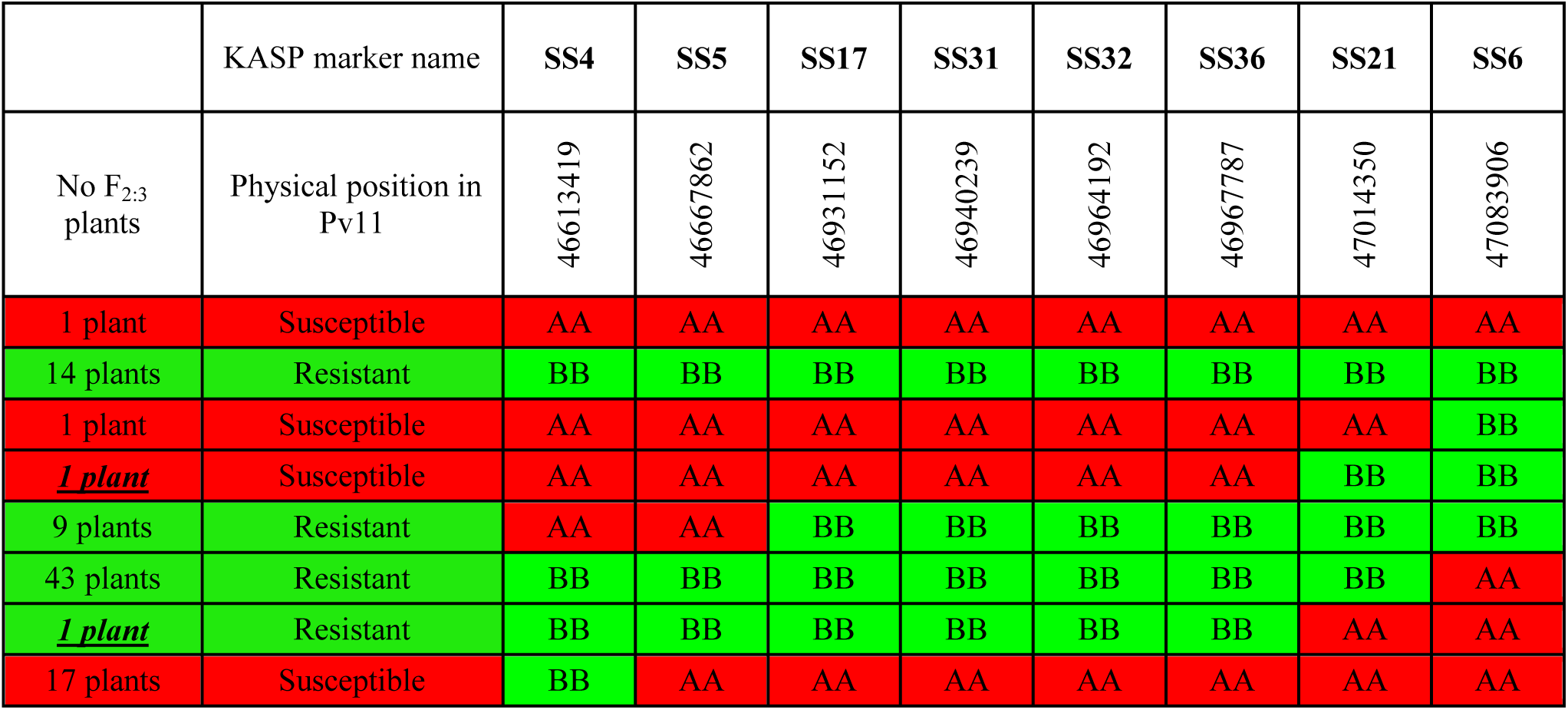
Genotypes at eight SNP loci and the reaction of 87 F_3_ plants with recombination events to race 53 of *Uromyces appendiculatus* in a Pinto 114 × Aurora population. The two F_3_ plants underlined had the same recombination breakpoint, but opposite phenotypes (AA = Pinto 114 allele; BB = Aurora allele) indicating the location of *Ur-3* locus.

Subsequent genotyping of the 129 F_2_ plants from the Pinto 114 × Aurora cross using KASP SS36 and KASP marker SS68, which was targeting the *Ur-3* haplotype and only~ 200bp downstream from SS36, showed that these markers were linked to the *Ur-3* rust resistance gene, with no recombination observed between bean rust phenotype and genotype (Table S2). The KASP marker SS68 effectively differentiated homozygous resistants, homozygous susceptibles, and heterozygous plants (Figure 2).

**Figure 2.**
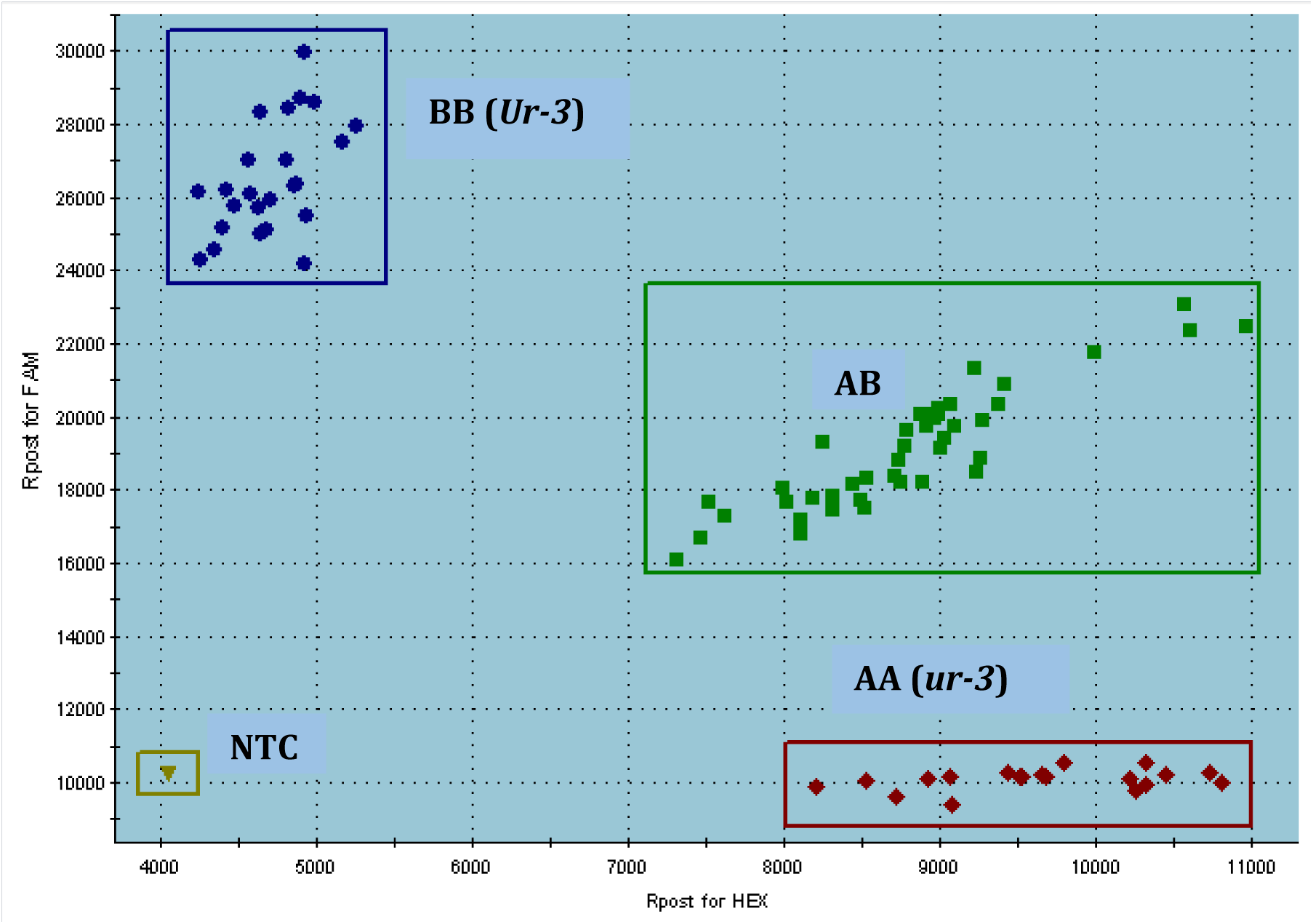
SNP graph of SS68 assay for the F_2_ plants from Pinto 114 × Aurora cross (BB = Aurora alleles; AA = Pinto 114 alleles; AB = heterozygous alleles, NTC= non-target control.

### Validation of KASP marker SS68 linked to *Ur-3* gene

We used the SS68 KASP marker to genotype a panel of 130 common bean cultivars and varieties that included dry and snap beans. Some of these common beans contained the *Ur-3* gene alone while others had *Ur-3* in combination with other rust resistance genes. In addition, other cultivars had single or combinations of the other ten rust resistance genes in common bean. The results of this validation showed that SS68 was highly accurate for the identification of the *Ur-3* locus (Table 7). No false positives or false negatives were observed when comparing the genotypic (evaluation with SS68 marker) and phenotypic (reaction to race 53) evaluations of these cultivars.

**Table 7.**
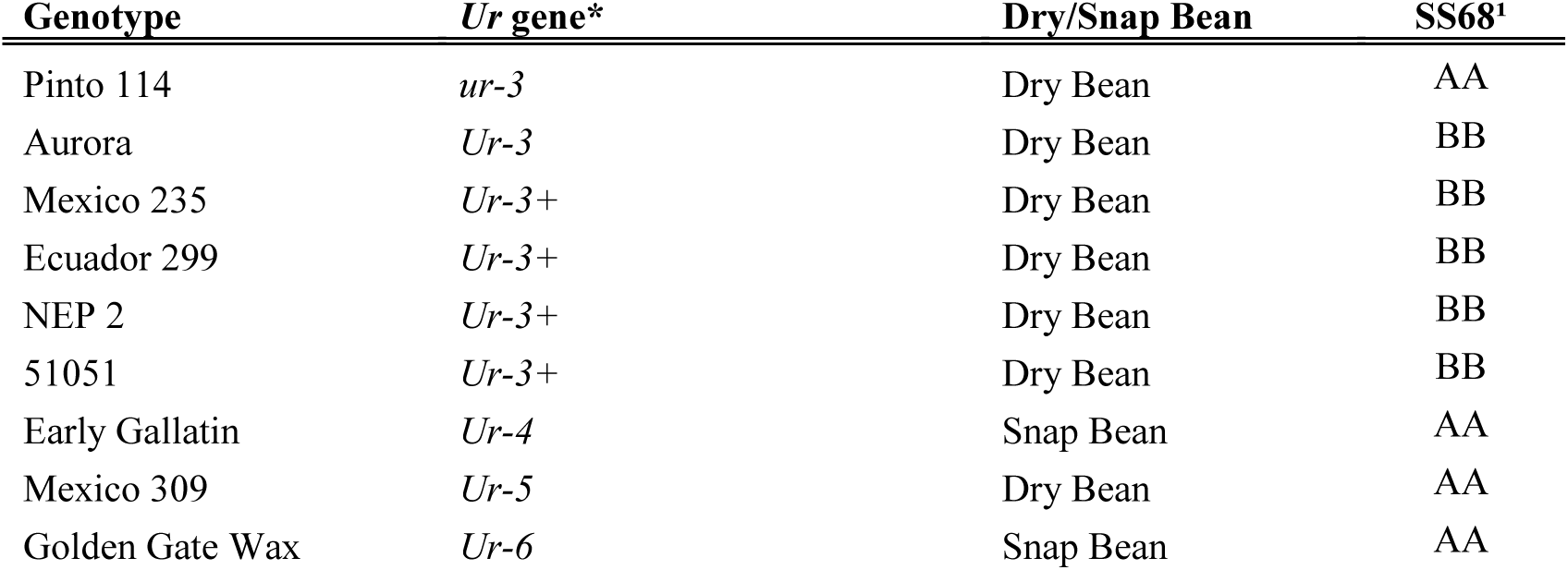
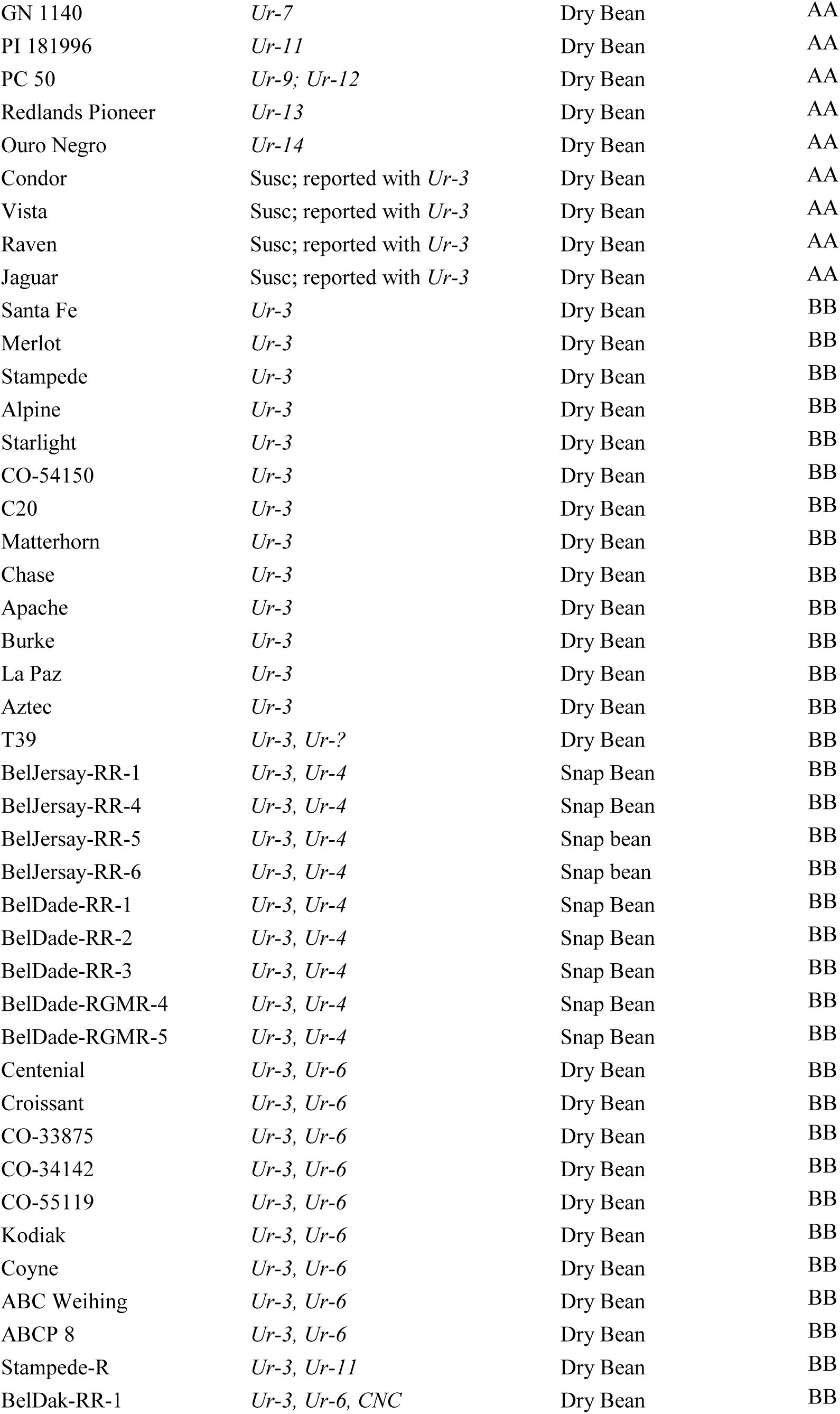
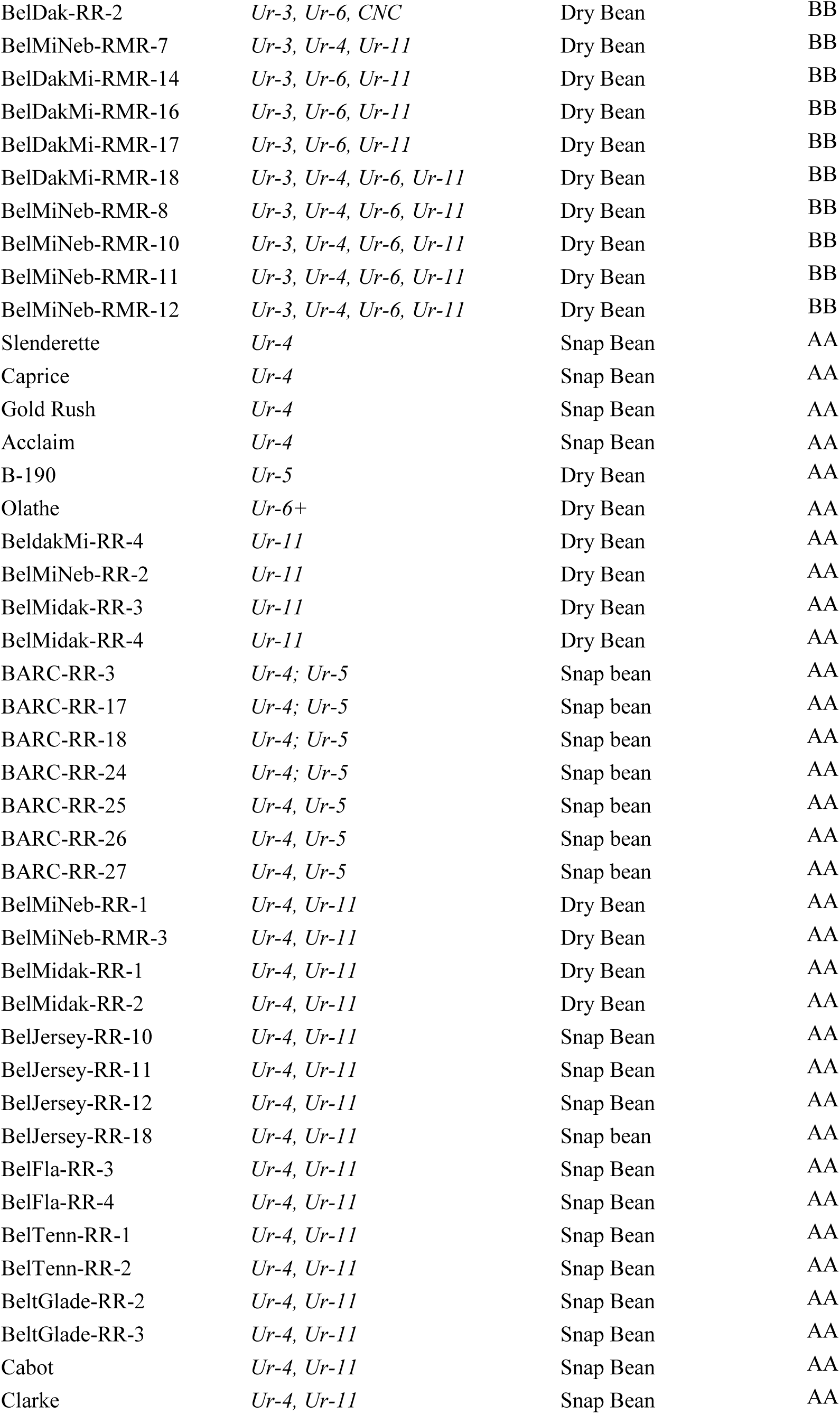
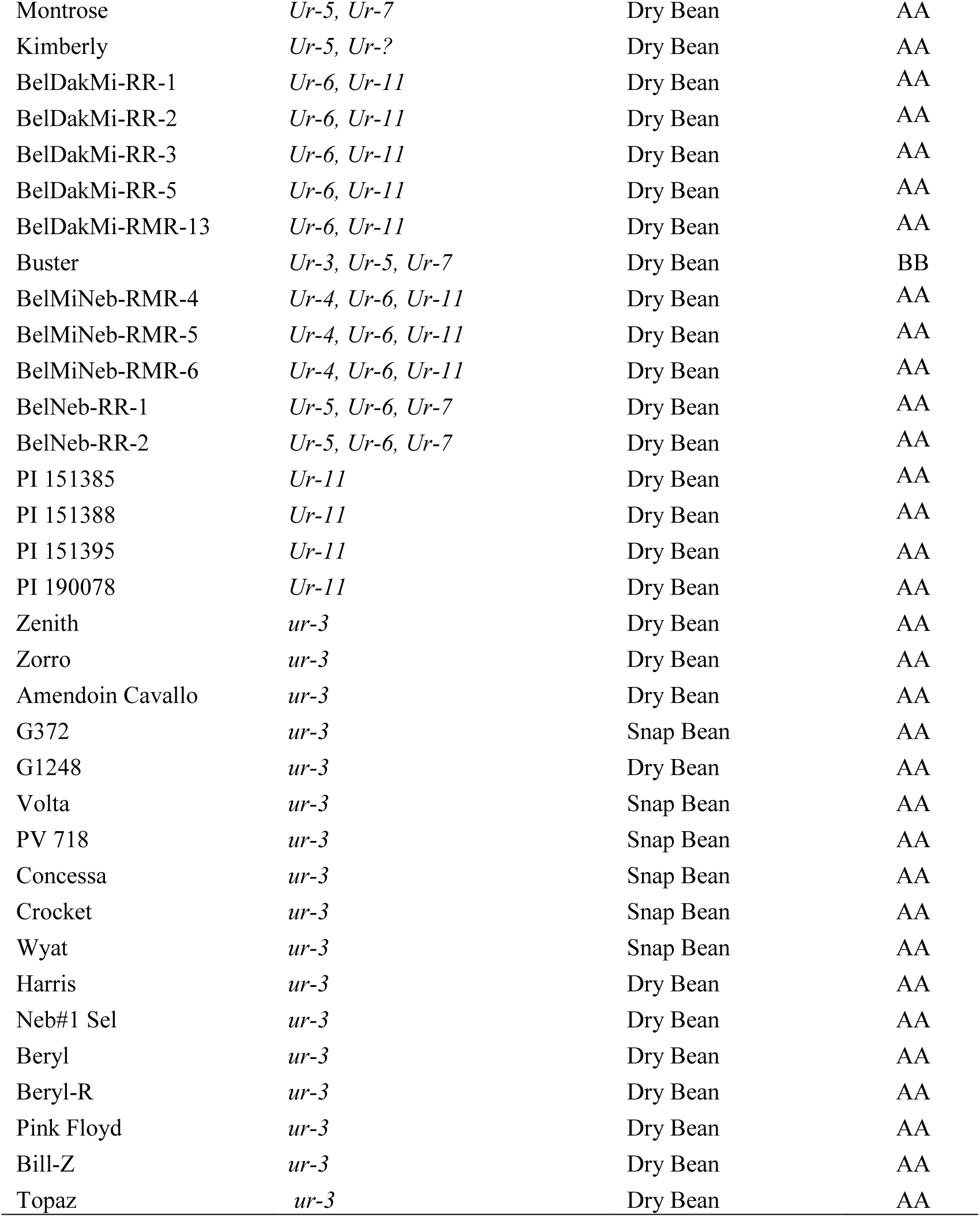
Validation of the single nucleotide polymorphism (SNP) marker associated to *Ur-3* locus on Pv11. Marker SS68 was tested in a common bean panel containing Andean and Mesoamerican cultivars with or without *Ur-3* gene representing most of the market classes. *=*Ur* gene identified based on phenotypic characterization using multiple races of *U. appendiculatus*.^1^ = allele score generated by KASP marker SS68 described in this study. CNC=Compuesto Negro de Chimaltenango, *Ur-?* denotes unknown rust resistant gene. Susc = Susceptible based on phenotype reaction to race 53 of *U. appendiculatus*.

| Genotype | *Ur* gene* | Dry/Snap Bean | SS68<sup>1</sup> |
| --- | --- | --- | --- |
| Pinto 114 | *ur-3* | Dry Bean | AA |
| Aurora | *Ur-3* | Dry Bean | BB |
| Mexico 235 | *Ur-3+* | Dry Bean | BB |
| Ecuador 299 | *Ur-3+* | Dry Bean | BB |
| NEP 2 | *Ur-3+* | Dry Bean | BB |
| 51051 | *Ur-3+* | Dry Bean | BB |
| Early Gallatin | *Ur-4* | Snap Bean | AA |
| Mexico 309 | *Ur-5* | Dry Bean | AA |
| Golden Gate Wax | *Ur-6* | Snap Bean | AA |
| GN 1140 | *Ur-7* | Dry Bean | AA |
| PI 181996 | *Ur-11* | Dry Bean | AA |
| PC 50 | *Ur-9; Ur-12* | Dry Bean | AA |
| Redlands Pioneer | *Ur-13* | Dry Bean | AA |
| Ouro Negro | *Ur-14* | Dry Bean | AA |
| Condor | Susc; reported with *Ur-3* | Dry Bean | AA |
| Vista | Susc; reported with *Ur-3* | Dry Bean | AA |
| Raven | Susc; reported with *Ur-3* | Dry Bean | AA |
| Jaguar | Susc; reported with *Ur-3* | Dry Bean | AA |
| Santa Fe | *Ur-3* | Dry Bean | BB |
| Merlot | *Ur-3* | Dry Bean | BB |
| Stampede | *Ur-3* | Dry Bean | BB |
| Alpine | *Ur-3* | Dry Bean | BB |
| Starlight | *Ur-3* | Dry Bean | BB |
| CO-54150 | *Ur-3* | Dry Bean | BB |
| C20 | *Ur-3* | Dry Bean | BB |
| Matterhorn | *Ur-3* | Dry Bean | BB |
| Chase | *Ur-3* | Dry Bean | BB |
| Apache | *Ur-3* | Dry Bean | BB |
| Burke | *Ur-3* | Dry Bean | BB |
| La Paz | *Ur-3* | Dry Bean | BB |
| Aztec | *Ur-3* | Dry Bean | BB |
| T39 | *Ur-3, Ur-?* | Dry Bean | BB |
| BelJersay-RR-1 | *Ur-3, Ur-4* | Snap Bean | BB |
| BelJersay-RR-4 | *Ur-3, Ur-4* | Snap Bean | BB |
| BelJersay-RR-5 | *Ur-3, Ur-4* | Snap bean | BB |
| BelJersay-RR-6 | *Ur-3, Ur-4* | Snap bean | BB |
| BelDade-RR-1 | *Ur-3, Ur-4* | Snap Bean | BB |
| BelDade-RR-2 | *Ur-3, Ur-4* | Snap Bean | BB |
| BelDade-RR-3 | *Ur-3, Ur-4* | Snap Bean | BB |
| BelDade-RGMR-4 | *Ur-3, Ur-4* | Snap Bean | BB |
| BelDade-RGMR-5 | *Ur-3, Ur-4* | Snap Bean | BB |
| Centenial | *Ur-3, Ur-6* | Dry Bean | BB |
| Croissant | *Ur-3, Ur-6* | Dry Bean | BB |
| CO-33875 | *Ur-3, Ur-6* | Dry Bean | BB |
| CO-34142 | *Ur-3, Ur-6* | Dry Bean | BB |
| CO-55119 | *Ur-3, Ur-6* | Dry Bean | BB |
| Kodiak | *Ur-3, Ur-6* | Dry Bean | BB |
| Coyne | *Ur-3, Ur-6* | Dry Bean | BB |
| ABC Weihing | *Ur-3, Ur-6* | Dry Bean | BB |
| ABCP 8 | *Ur-3, Ur-6* | Dry Bean | BB |
| Stampede-R | *Ur-3, Ur-11* | Dry Bean | BB |
| BelDak-RR-1 | *Ur-3, Ur-6, CNC* | Dry Bean | BB |
| BelDak-RR-2 | *Ur-3, Ur-6, CNC* | Dry Bean | BB |
| BelMiNeb-RMR-7 | *Ur-3, Ur-4, Ur-11* | Dry Bean | BB |
| BelDakMi-RMR-14 | *Ur-3, Ur-6, Ur-11* | Dry Bean | BB |
| BelDakMi-RMR-16 | *Ur-3, Ur-6, Ur-11* | Dry Bean | BB |
| BelDakMi-RMR-17 | *Ur-3, Ur-6, Ur-11* | Dry Bean | BB |
| BelDakMi-RMR-18 | *Ur-3, Ur-4, Ur-6, Ur-11* | Dry Bean | BB |
| BelMiNeb-RMR-8 | *Ur-3, Ur-4, Ur-6, Ur-11* | Dry Bean | BB |
| BelMiNeb-RMR-10 | *Ur-3, Ur-4, Ur-6, Ur-11* | Dry Bean | BB |
| BelMiNeb-RMR-11 | *Ur-3, Ur-4, Ur-6, Ur-11* | Dry Bean | BB |
| BelMiNeb-RMR-12 | *Ur-3, Ur-4, Ur-6, Ur-11* | Dry Bean | BB |
| Slenderette | *Ur-4* | Snap Bean | AA |
| Caprice | *Ur-4* | Snap Bean | AA |
| Gold Rush | *Ur-4* | Snap Bean | AA |
| Acclaim | *Ur-4* | Snap Bean | AA |
| B-190 | *Ur-5* | Dry Bean | AA |
| Olathe | *Ur-6+* | Dry Bean | AA |
| BeldakMi-RR-4 | *Ur-11* | Dry Bean | AA |
| BelMiNeb-RR-2 | *Ur-11* | Dry Bean | AA |
| BelMidak-RR-3 | *Ur-11* | Dry Bean | AA |
| BelMidak-RR-4 | *Ur-11* | Dry Bean | AA |
| BARC-RR-3 | *Ur-4; Ur-5* | Snap bean | AA |
| BARC-RR-17 | *Ur-4; Ur-5* | Snap bean | AA |
| BARC-RR-18 | *Ur-4; Ur-5* | Snap bean | AA |
| BARC-RR-24 | *Ur-4; Ur-5* | Snap bean | AA |
| BARC-RR-25 | *Ur-4, Ur-5* | Snap bean | AA |
| BARC-RR-26 | *Ur-4, Ur-5* | Snap bean | AA |
| BARC-RR-27 | *Ur-4, Ur-5* | Snap bean | AA |
| BelMiNeb-RR-1 | *Ur-4, Ur-11* | Dry Bean | AA |
| BelMiNeb-RMR-3 | *Ur-4, Ur-11* | Dry Bean | AA |
| BelMidak-RR-1 | *Ur-4, Ur-11* | Dry Bean | AA |
| BelMidak-RR-2 | *Ur-4, Ur-11* | Dry Bean | AA |
| BelJersey-RR-10 | *Ur-4, Ur-11* | Snap Bean | AA |
| BelJersey-RR-11 | *Ur-4, Ur-11* | Snap Bean | AA |
| BelJersey-RR-12 | *Ur-4, Ur-11* | Snap Bean | AA |
| BelJersey-RR-18 | *Ur-4, Ur-11* | Snap bean | AA |
| BelFla-RR-3 | *Ur-4, Ur-11* | Snap Bean | AA |
| BelFla-RR-4 | *Ur-4, Ur-11* | Snap Bean | AA |
| BelTenn-RR-1 | *Ur-4, Ur-11* | Snap Bean | AA |
| BelTenn-RR-2 | *Ur-4, Ur-11* | Snap Bean | AA |
| BeltGlade-RR-2 | *Ur-4, Ur-11* | Snap Bean | AA |
| BeltGlade-RR-3 | *Ur-4, Ur-11* | Snap Bean | AA |
| Cabot | *Ur-4, Ur-11* | Snap Bean | AA |
| Clarke | *Ur-4, Ur-11* | Snap Bean | AA |
| Montrose | *Ur-5, Ur-7* | Dry Bean | AA |
| Kimberly | *Ur-5, Ur-?* | Dry Bean | AA |
| BelDakMi-RR-1 | *Ur-6, Ur-11* | Dry Bean | AA |
| BelDakMi-RR-2 | *Ur-6, Ur-11* | Dry Bean | AA |
| BelDakMi-RR-3 | *Ur-6, Ur-11* | Dry Bean | AA |
| BelDakMi-RR-5 | *Ur-6, Ur-11* | Dry Bean | AA |
| BelDakMi-RMR-13 | *Ur-6, Ur-11* | Dry Bean | AA |
| Buster | *Ur-3, Ur-5, Ur-7* | Dry Bean | BB |
| BelMiNeb-RMR-4 | *Ur-4, Ur-6, Ur-11* | Dry Bean | AA |
| BelMiNeb-RMR-5 | *Ur-4, Ur-6, Ur-11* | Dry Bean | AA |
| BelMiNeb-RMR-6 | *Ur-4, Ur-6, Ur-11* | Dry Bean | AA |
| BelNeb-RR-1 | *Ur-5, Ur-6, Ur-7* | Dry Bean | AA |
| BelNeb-RR-2 | *Ur-5, Ur-6, Ur-7* | Dry Bean | AA |
| PI 151385 | *Ur-11* | Dry Bean | AA |
| PI 151388 | *Ur-11* | Dry Bean | AA |
| PI 151395 | *Ur-11* | Dry Bean | AA |
| PI 190078 | *Ur-11* | Dry Bean | AA |
| Zenith | *ur-3* | Dry Bean | AA |
| Zorro | *ur-3* | Dry Bean | AA |
| Amendoin Cavallo | *ur-3* | Dry Bean | AA |
| G372 | *ur-3* | Snap Bean | AA |
| G1248 | *ur-3* | Dry Bean | AA |
| Volta | *ur-3* | Snap Bean | AA |
| PV 718 | *ur-3* | Snap Bean | AA |
| Concessa | *ur-3* | Snap Bean | AA |
| Crocket | *ur-3* | Snap Bean | AA |
| Wyat | *ur-3* | Snap Bean | AA |
| Harris | *ur-3* | Dry Bean | AA |
| Neb#1 Sel | *ur-3* | Dry Bean | AA |
| Beryl | *ur-3* | Dry Bean | AA |
| Beryl-R | *ur-3* | Dry Bean | AA |
| Pink Floyd | *ur-3* | Dry Bean | AA |
| Bill-Z | *ur-3* | Dry Bean | AA |
| Topaz | *ur-3* | Dry Bean | AA |

### The *Ur-3* locus contains six candidate genes

The genomic region delimited by markers SS36 and SS21 defined as the *Ur-3* locus, contained six candidate genes according to Phytozome.net database for *P vulgaris* assembly V1.0. The names of these genes are: Phvul.011G193100, Phvul.011G193200, Phvul.011G193300, Phvul.011G193400, Phvul.011G193500 and Phvul.011G193600. Three of these *Ur-3* genes (Phvul.011G193100, Phvul.011G193500 and Phvul.011G193600) are classified as containing NB-ARC domains and leucine rich repeat (LRR) regions. Genes Phvul.011G193200 and Phvul.011G193400 are annotated as serine/threonine kinases, and Phvul.011G193300 is a tyrosine kinase with salt/stress response-related and antifungal function. All these candidate genes except Phvul.011G193600 were highly expressed in common bean leaves according to the expression level experiments recorded in the JGI genome browser for *P. vulgaris*.

## Discussion

### Development of accurate SNP markers linked to *Ur-3* locus

The historically important *Ur-3* gene confers resistance to the pathogen that causes the rust disease of common bean. The effective incorporation of *Ur-3* into dry and snap beans using molecular markers has been limited by the inaccuracy of the markers linked to this gene (Stavely 2000). The reported RAPD (OK14_620_) and the SCAR (SK14) markers linked to *Ur-3* produced false-positive results (Haley et al 1994; Nemchinova and Stavely 1998). More recently, we have used bulk segregant analysis, single nucleotide polymorphism assay, and whole genome sequencing to discover simple sequence repeat markers closely linked to the *Ur-3* and other disease resistance genes. However, even the use of closely linked BARCPVSSR14007, an SSR marker reported in this study that mapped at 0.2 cM from the *Ur-3* locus, resulted in more than 3% false positive results when this marker was used on the panel of 130 common bean lines and varieties (data not shown). As indicated earlier, the inability to find specific molecular markers linked to *Ur-3* may have been exacerbated by the presence of the *Ur-11* rust resistance gene that is closely linked to *Ur-3* on the terminal position of chromosome Pv11. Currently, the most reliable method to monitor for the presence of the *Ur-3* gene in dry and snap bean cultivars continues to be race 53 of *U. appendiculatus.* Race 53 is used as a phenotypic marker that effectively identifies common bean plants with *Ur-3* alone or in combination. (Pastor-Corrales 2002). However, phenotypic evaluations under greenhouse conditions are very laborious and time consuming (about 21 days). Moreover, due mostly, but not only, to the biotrophic condition of the rust pathogen, most breeders of dry and snap bean in the world do not have the option of using this methodology.

Given the importance of *Ur-3*, we determined to search for highly accurate molecular markers linked to *Ur-3*, using a fine mapping approach. We employed a variety of technologies that included phenotyping with specific races of the bean trust pathogen, bulk segregant analysis coupled with high-throughput SNP genotyping using the BARCBEAN6K_3 BeadChip, SSR and KASP marker development, and local association analysis using SNPs from previous whole genome shotgun sequencing efforts. In summary, these technologies permitted the identification of KASP marker SS68 that was highly accurate in identifying the presence of *Ur-3* in a panel of 130 common bean lines and cultivars that included dry and snap beans with and without the *Ur-3* gene. Marker SS68 was also tested on a mapping population of 184 F_2_ genotypes from the cross between Pinto 114 x Mexico 235 (*Ur-3*+). No recombination was observed between phenotype and the genotype in this study (data not shown). These results confirm the accuracy and utility of the KASP marker SS68 even when this marker is used on mapping populations in which the origin of the *Ur-3* gene is not the cultivar Aurora.

### Survey of the SS68 KASP marker in a common bean diversity panel

In this work we determined the potential utility of the KASP SNP marker SS68 in a panel of common bean cultivars carrying different rust resistance genes and in bean lines representing the major common bean market classes in the United States. Marker SS68 reliably identified cultivars containing *Ur-3*, independent of the gene pool (Andean or Mesoamerican), type of common bean (dry or snap), or market class of dry edible beans (pinto, great northern, navy, red kidney, black, and others). Additionally SS68 effectively distinguished common bean lines carrying *Ur-3* alone from lines combining the Ur-3 and *Ur-11* genes that are closely linked on Pv11, which could not be accomplished with previously reported markers linked to *Ur-3* (Table 7). Moreover because *Ur-3* gene is epistatic to *Ur-11*, it is difficult to combine these two genes using inoculations with races of the rust pathogen (Stavely, 2000). Thus, using marker SS68 to identify *Ur-3* when combined with *Ur-11* can avoid this hurdle.

### The *Ur-3* locus contains a cluster of R genes

Through haplotype analysis and KASP marker development, it was possible to determine a genomic region of 46,563 bp containing the *Ur-3* locus and delimited by markers SS36 and SS21 on Pv11. Six candidate genes were identified within this 46.5 Kb region in the *P. vulgaris* reference genome, obtained by sequencing the landrace G19833 of Andean origin. Among the six candidate genes, there were three genes with NB-ABC LRR domains. Proteins containing NB-LRR domain are known to be involved in plant resistance and activation of innate immune responses to various types of pathogens (Hammond-Kosack and Jones 1997; Jones and Dangl 2006). Similarly, protein kinases, also found in the 46.5 Kb region, are known to play a central role in signaling during pathogen recognition and the subsequent activation of plant defense mechanisms (Xue et al. 2015). The genomic region containing *Co-4* gene on chromosome Pv08, conferring resistance to *Colletotrichum lindemuthianum* in common bean, has been characterized and known to contain an open reading frame coding for a serine threonine kinase (Oblessuc et al. 2015) a type of protein which has also been identified in our studies. Additionally, serine threonine protein kinase constitutes candidate genes for controlling angular leaf spot resistance in the Andean landrace G 5686 (Keller et al. 2015). Whether the phenotype of the *Ur-3* locus is the result of the expression of one or more of the six candidates genes will be a matter of further investigation.

Sequence analysis of the Andean landrace G 19833 used to sequence the reference genome of common bean, which is susceptible to race 53, indicating that it does not have *Ur-3*, revealed that the 46.5 Kb genomic region containing the *Ur-3* locus, is highly duplicated (Figure S1) and it includes repetitive elements in the intergenic spaces. Additionally, this genomic region is AT-rich (33% vs. 16% for GC) which suggests that it is highly unstable. Sequence analysis comparing the Mesoamerican Aurora common bean and the Andean landrace G19833, will provide valuable insights into the structural differences and evolutionary history of the important *Ur-3* rust resistance locus.

## CONCLUSIONS

This study used a new approach to generate KASP SS68, the first highly accurate DNA marker linked to the *Ur-3* rust resistance gene in common bean. We fine-mapped a 46.5 kb genomic region in chromosome Pv11 present in Mesoamerican common bean cultivar Aurora. This was accomplished using the BARCBEAN6K_3 BeadChip, SSRs, KASP technology, and local association. The validation of this newly discovered SS68 marker on a panel of 130 common bean lines and varieties revealed that SS68 was highly accurate in identifying *Ur-3*. This marker will be of great value for common beans combining *Ur-3* with other Andean and Mesoamerican genes with broad-spectrum resistance to the highly variable bean rust pathogen. In addition, the utilization of the new KASP marker SS68 will reduce significantly the time and labor associated with the transfer of the *Ur-3* gene using inoculations of bean plants with specific races of the rust pathogen. The genomic region containing the *Ur-3* locus included six genes annotated in the reference genome of *P. vulgaris*. These genes are likely candidates for the *Ur-3* rust resistance gene. Gene expression analysis of these candidate genes and functional approaches will enhance our understanding of the mechanisms underlying the reaction of *P. vulgaris* to *Uromyces appendiculatus*.

## Acknowledgments

This work was supported, in part, by funding from the Norman Borlaug Commemorative Research Initiative of the US Agency for International Development, project number 0210-22310-004-96R and by the US Department of Agriculture, Agricultural Research Services Project Number 8042-22000-286-00D (Pastor-Corrales). The contents of this publication do not necessarily reflect the views or policies of the US Department of Agriculture, nor does mention of trade names, commercial products, or organizations imply endorsement by the US Government. This research was also financially supported by the National Council for Scientific and Technological Development (CNPQ) and the Coordination for the Improvement of Higher Education Personnel (CAPES). The authors also thank Rob Parry and Chris Pooley for their assistance with sequence analysis by installing computer hardware and software.

## Supplementary Information

**Figure S1.**
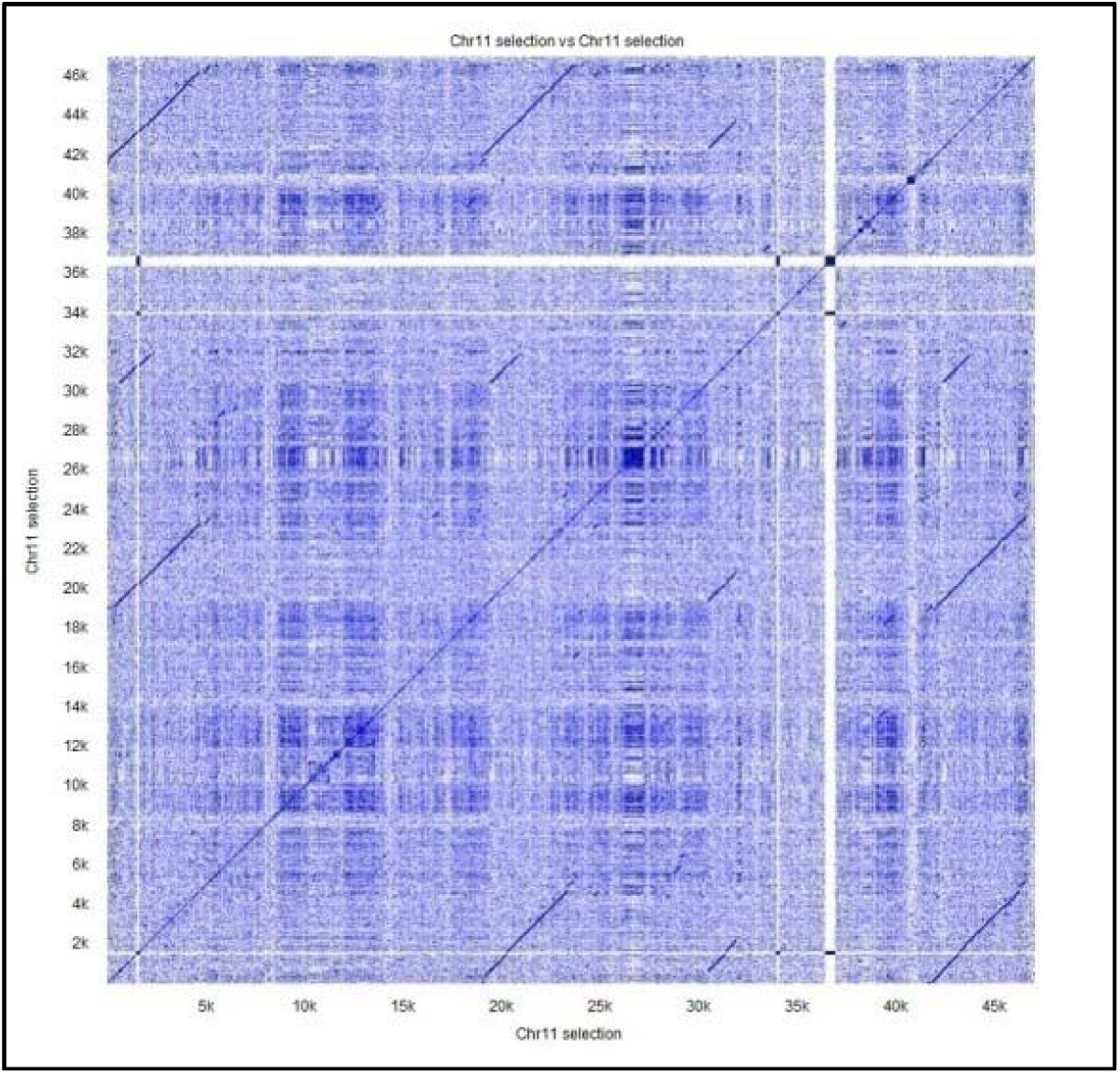
Dot blot comparison of the *Ur-3* locus comprising the 46.5 kb region identified through fine mapping in this study. Sequence used for the analysis is from the reference genome G19833. Small solid lines above or below the diagonal line suggest the presence of duplicated or highly similar sequences within the region. Analysis done using CLC Genomics Workbench following manual instructions.

